# Long-term consolidation switches goal proximity coding from hippocampus to retrosplenial cortex

**DOI:** 10.1101/167882

**Authors:** E. Zita Patai, Amir-Homayoun Javadi, Jason D. Ozubko, Andrew O’Callaghan, Shuman Ji, Jessica Robin, Cheryl Grady, Gordon Winocur, Shayna R. Rosenbaum, Morris Moscovitch, Hugo J. Spiers

**Author notes:** These authors contributed equally (E. Zita Patai, Amir-Homayoun Javadi). Corresponding Author: Hugo Spiers, Institute of Behavioural Neuroscience, UCL Department of Experimental Psychology, 26 Bedford Way, London, WC1H 0AP, UK.

## Abstract

Recent research indicates the hippocampus may code the distance to the goal during navigation of newly learned environments. It is unclear however, whether this also pertains to highly familiar environments where extensive systems-level consolidation is thought to have transformed mnemonic representations. Here we recorded fMRI while University College London and Imperial College London students navigated virtual simulations of their own familiar campus (> 2 years of exposure) and the other campus learned days before scanning. Posterior hippocampal activity tracked the proximity to the goal in the newly learned campus, but not in the familiar campus. By contrast retrosplenial cortex tracked the distance to the goal in the familiar campus, but not in the recently learned campus. These responses were abolished when participants were guided to their goal by external cues. These results open new avenues of research on navigation and consolidation of spatial information and help advance models of how neural circuits support navigation in novel and highly familiar environments.

**Significance Statement:** Historically, research on the hippocampal formation has focused on its role in long-term memory and navigation – often in isolation. No study to date has directly compared realistic navigation within familiar with recently learned environments, nor has it been explored how the neural substrates, along with computational codes, may change. In this study, we show for the first time, a shift from hippocampal to cortical coding of distance to a goal during active navigation. This study bridges the gap between memory consolidation and navigation, and paves the way for more functional and realistic understanding of the hippocampus.

## Introduction

Understanding how the brain consolidates memories is a central question in neuroscience (1). Historically, research has focused on contextual and recognition memory in rodents and primates (2–4) and episodic memory in humans (5–7). Despite substantial interest in the neural circuits that support navigation, little research has explored consolidation of spatial memories and representations of environments learned months or years ago (“familiar” environments) (8–10). This dearth of research is particularly surprising considering that current theories disagree about the contribution of the hippocampus to processing spatial representations over time: the standard consolidation theory (SCT) argues that initially the hippocampus is involved in processing the spatial memories and representations, and that over time the representations in neocortical regions are strengthened, reducing the demand on the hippocampus (11, 12). By contrast multiple trace theory (MTT), and its offspring Trace Transformation Theory, argue that for detailed spatial memories and representations the hippocampus is always involved, or more specifically that an episodic hippocampal trace will exist in addition to the schematized representation in the cortex that can be activated depending on task requirements (9, 13–16).

Neuropsychological evidence indicates that complex spatial memories acquired years in the past can become independent of the hippocampus (17–21) consistent with SCT. In several cases, however, hippocampal damage does appear to lead to impaired spatial memories for some detailed aspects of the environment (Herdman, Calarco, Moscovitch, Hirshhorn, & Rosenbaum, 2015; Maguire, Nannery, & Spiers, 2006; Rosenbaum et al., 2000), consistent with MTT. Insight from functional magnetic resonance imaging (fMRI) research has been highly limited. Only two prior fMRI experiments have examined navigation of familiar environments. One study involving London taxi drivers navigating a virtual simulation of London (UK) reported that the hippocampus is engaged at the start of navigating this highly familiar environment (22). The other study involved residents of Toronto mentally navigating this city and found no increased activity in the hippocampus (23). Crucially, however, neither study directly compared navigation in familiar with recently learned environments. Although there is evidence from fMRI that the hippocampus is less implicated in mentally navigating in familiar environments (Hirshhorn, Grady, Rosenbaum, Winocur, & Moscovitch, 2012), exactly what the contribution of the hippocampus and other structures is and the nature of their computations is not known. Additionally, as these studies have relied on static, mental navigation, they may not engage hippocampal activity as would active, dynamic navigation in a virtual reality environment. Thus, for now, we cannot rule out whether the differences in hippocampal findings relate to the demands of navigating different cities, with London placing greater demands on mental simulation of future familiar routes than Toronto, or whether the structure of the environment is key to these differences (10).

One question not yet addressed is whether long-term consolidation changes the spatial information processed by brain regions during the navigation of an environment. In recently learned environments, the hippocampus has been shown to encode the distance to the goal (Balaguer, Spiers, Hassabis, & Summerfield, 2016; Chrastil, Sherrill, Hasselmo, & Stern, 2015; Howard et al., 2014; Sherrill et al., 2013; Spiers & Maguire, 2007a; Viard, Doeller, Hartley, Bird, & Burgess, 2011). It is unknown whether this is also the case in highly familiar environments. Current models argue that through systems consolidation, neocortical regions (such as parahippocampal cortex or anterior cingulate cortex) may come to code such information in extensively learned environments (10). Candidate neocortical regions include the retrosplenial cortex (32, 33), parahippocampal cortex (Bohbot et al., 2015; Rosenbaum et al., 2004, for extensive reviews see Epstein, 2008; Miller et al., 2014; Ranganath & Ritchey, 2012; Spiers & Maguire, 2007b), and the anterior cingulate cortex (38). It is also possible that the involvement of brain regions will vary depending on how individuals plan their route or use certain strategies, with MTT/TTT predicting that the hippocampus, and in particular the posterior hippocampus, would play a more important role when perceptually detailed processing is required (39).

Here we combined fMRI and a virtual simulation of two university campuses to examine the brain regions coding the distance to the goal in highly familiar and recently learned environments within the same scan session. Students from two London universities (University College London and Imperial College London) navigated each campus, with the one they were not attending made familiar via training material and a walking tour days before the fMRI session. During the scanning session they engaged in active navigation towards goal locations and subsequently reported on when they planned their routes. We aimed to test whether a) the hippocampus codes distances to the goal, b) this would depend on which environment they were in, and if this code may be represented elsewhere in the brain after extended exposure, and c) how these representations were related to route planning.

## Results

Our experimental fMRI task was adapted from Howard et al. (2014), in which participants (n = 25) were presented with a goal location in a virtual simulation of the environment and required to travel along the streets (Travel Periods) and make path choices prior to street junctions (Decision Points), see Supplemental Methods and Figure 1. In matched control routes participants were instructed which street to select during navigation to goal locations. Our task differed in that there were two environments to navigate: a familiar campus (‘familiar’) and a new campus (‘recent’). Participants were exposed via an intensive *in situ* training tour of both campuses in the immediate preceding days before scanning, ensuring that the only difference between the two environments was the long-term (> two year) prior knowledge of one of them. Behavioural results revealed that participants were able to orient themselves and make correct decisions at Decision Points in both environments, albeit better so in their familiar campus (Familiar: M = 89.9%, SD = 11%; Recent: M = 80.4%, SD = 13%, paired sample t-test: t(24)=3.6, p=0.002). Participants were also faster to respond in their familiar environment (Familiar = 1.19 ± 0.4s, Recent = 1.39 ± 0.5s, t(24)=4.7, p<.001), and reaction times scaled with distance to the goal along the future path (across subjects and junctions, both environments r(124)>.24, p<0.01). After scanning participants completed a debrief session, which consisted of filling out a questionnaire regarding their navigational strategies, as well as a behavioural version of the task, in which they indicated whether they had engaged in planning during Decision Points. This was only done for the active navigation routes (see Supplemental Methods & Results for more details).

**Figure 1:**
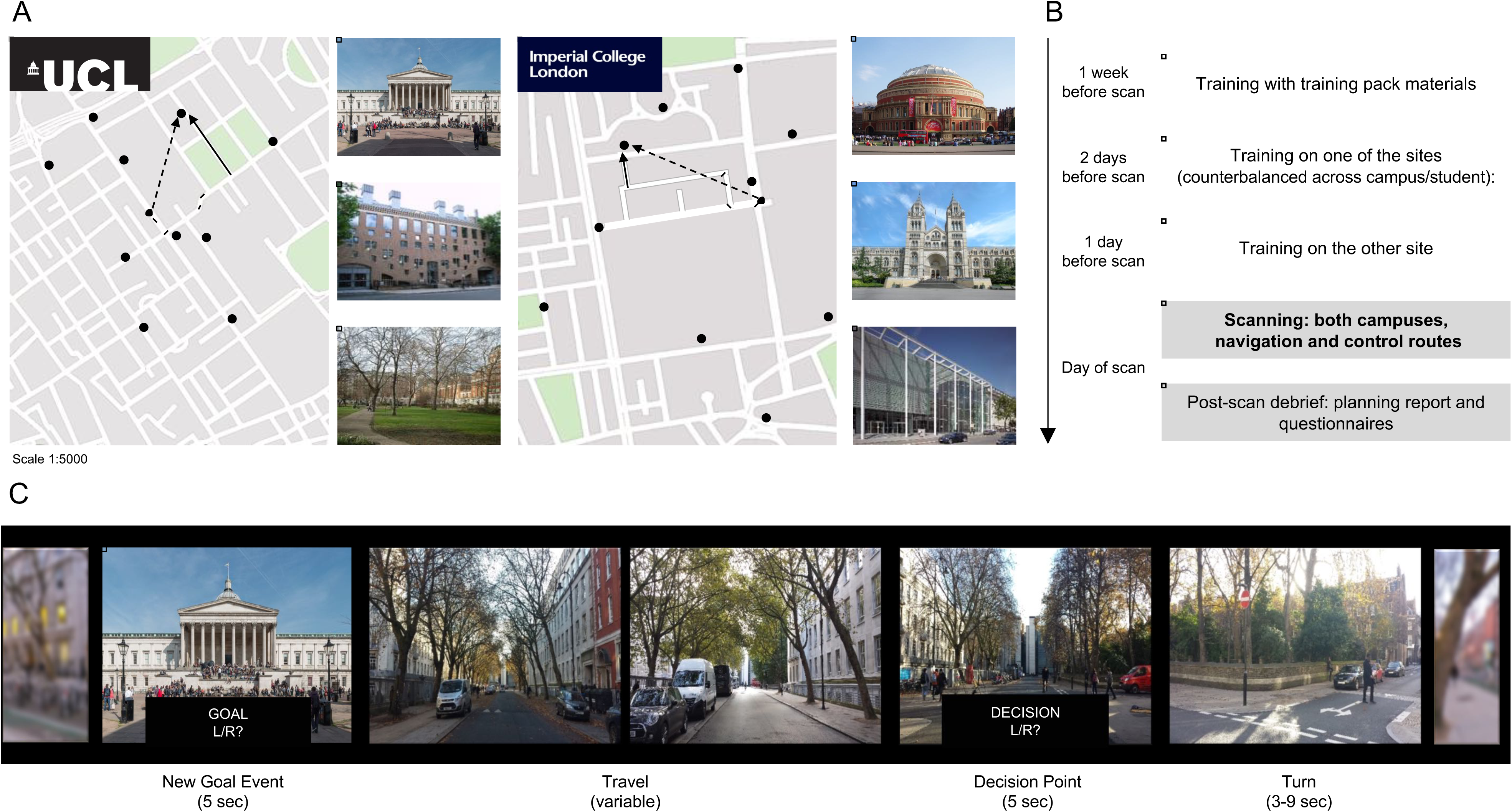
Campus layouts and training protocol. A) Participants navigated around either UCL (left) or Imperial College campus (right), finding their way to various goal locations (examples depicted to the right of the maps – note not all goal locations were necessarily famous). The maps show all potential goal locations marked with black circles. The open circle represents an example participant location, the solid and dashed black lines indicate the optimal path to be taken (Path Distance), and Euclidian distance (“as the crow flies”), respectively, to the current goal location. B) The training protocol consisted of two stages. First, a self-guided learning of landmarks and campus layouts with printed training materials. Two days before the fMRI scan participants were taken on an intensive guided tour of one of the campuses, and the next day, they were taken around the other campus. The order was counterbalanced across subjects and could start at either their home campus or the new one. On the day of scanning, participants completed routes in both campuses, in both active navigation and a control condition that involved following directions. During the debrief session, they filled out questionnaires regarding navigation strategies, and performed a behavioural task where they indicated where along the route they engaged in planning while navigating. C) Excerpt from a “navigation” route at UCL. At the start of every route, participants were shown the street they are on, as well as their facing direction. Next they were given a New Goal Location, and subsequently asked to indicate the general direction of that landmark. They travelled down the road until they reach a junction (the duration and number of images of this was variable depending on the length of the street), at which point they were asked to indicate the correct turn to take towards the goal (Decision Point). Here, the program automatically advanced on a pre-defined route, which may have been correct, or it may have been an unplanned detour (occurring less frequently). After a variable interval, a new street was entered and the participant was informed again of their location and facing direction. Note the jittered interval after Decision Points is to allow for separating signals relating to Decision Points and Turns (or Detours). Analysis of fMRI data was constrained to the Travel and Decision Point periods. For fMRI analysis, Travel periods were taken as the mid-point between two events, and modelled as a punctate event. For simplicity, the above figure only depicts part of a route, and Travel is shown as two frames, but could range between 3–48 frames (on average: mean 31±18).

## fMRI Analysis

A categorical analysis of different event types and task blocks revealed more activity in lateral and medial parietal areas in familiar environments when actively navigating (“navigation” condition) compared to just following along a route (“control” condition), whereas recent environments did not show such a distinction (Figure S1). There were no clear differences when comparing familiar to recent navigation (Figure S2, and Supplementary Results). In order to examine how the neural responses relate to metrics in computational models we interrogated our fMRI data with parameters related to the distance and direction to the goal. We explored how path distance to the goal (Figure 1A) was correlated with brain activity during events sampled during Travel Periods and at Decision Points. We focused on these events because we were able to sample >20 events per condition (familiar/recent, navigation/control).

### Hippocampus and retrosplenial cortex track path distance to the goal during travel periods in recent and familiar environments, respectively

During Travel Periods when navigating the recently learned environment, we found a significant negative correlation with the distance to the goal in the right mid-posterior hippocampus (Figure 2A, Table S4), indicating that hippocampal activity increased with proximity to the goal. This effect was absent in the familiar environment and the control routes (Figure 2A, right panel). Moreover, right posterior hippocampal activity was significantly more correlated with proximity to the goal during navigation of the recently learned environment than the other conditions combined (Figure 2C, Table S4). Although the hippocampus did not appear to track the distance to the goal in the highly familiar environment, we found that the retrosplenial cortex did. During Travel Periods of navigation routes in the familiar environment we observed a significant positive correlation with distance in the retrosplenial cortex (Figure 2B, Table S4), indicating the activity was greatest when participants were farthest from their goal. This response was absent in the recent environment and during the control conditions (Figure 2B, right panel). Retrosplenial activity was significantly more correlated with the distance to the goal in familiar navigation routes than the other conditions combined (Figure 2D, Table S4). Even at liberal thresholds, activity in our other predicted ROIs (anterior cingulate cortex, caudate and parahippocampal cortices) was not significantly correlated with distance to the goal during Travel Periods in either recent or familiar environments (Figure S3).

**Figure 2.**
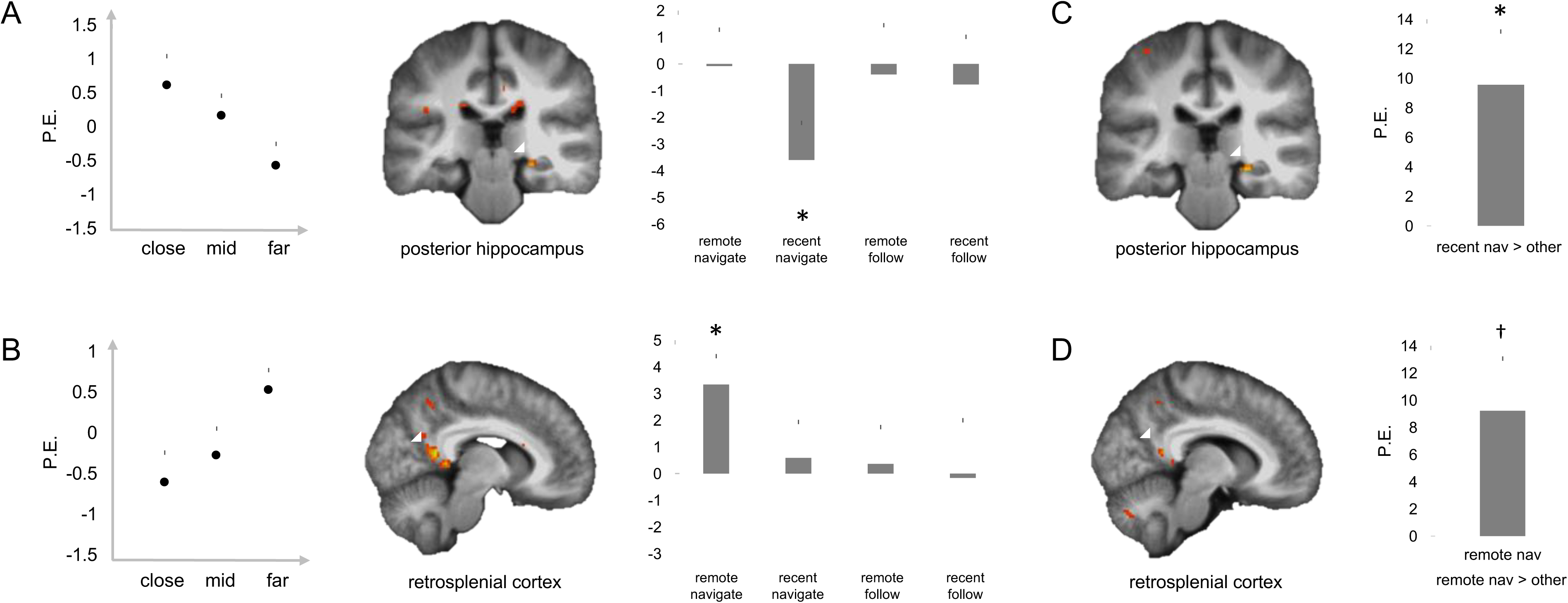
Travel distance in remote and recent environments is coded in different brain areas. A) During Travel in recent environments, there was a significant negative correlation with path distance, such that there was higher BOLD activity in the hippocampus when participants were closer to the goal location. B) During Travel in remote environments, there was a significant positive correlation with path distance, such that there was higher BOLD activity in the retrosplenial cortex when participants were further away from the goal location. In each plot, from left to right: parameter estimates (PE) extracted from a categorical model (binned by distance), the BOLD activity for the relevant condition, and the PE from the peak voxel in the ROI for each condition. All effects survive small-volume correction, including when ED (Euclidian distance) is added to the models. C/D) Show the data when the GLM included weighted regressors for the effects seen in A/B, respectively. For example, in C, the contrast was 1 −3 1 1, testing for an overall effect of correlation with path distance during Travel, for the recent navigation condition compared to all other conditions. The hippocampal and retrosplenial effects from A and B, respectively, are replicated indicating that the correlations with path distance are selective to these areas. *=p<0.05 SVC, †p<0.1 SVC

In our previous study (Howard et al., 2014) we found that during the Travel Period hippocampal activity was positively, rather than negatively, correlated with the distance to the goal. Because the previous study used a much smaller sized environment (200-400m route lengths) than the current one (200-1000m route lengths), we examined the impact of down-sampling the range distances in our analysis, excluding those over 900m (accounting for the top 25% of distances, see Figure S4A and Supplementary Methods). This analysis abolished the correlations between distance and activity in hippocampus and retrosplenial cortex, indicating that the larger distances were important in driving the relationship between distance and activity. Notably, the correlations, however, were not abolished when we down-sampled the data by removing the same number of events, but randomly sampled across the whole distance range (Figure S4A, lower panel). Though in this study path distance and Euclidian distance were highly correlated (Table S1), we ran a control model including both spatial parameters, and found both the hippocampal and retrosplenial results remained significant (Table S4).

### Hippocampal activity is positively correlated with the distance to the goal at decision points

At Decision Points during the navigation of the recently learned environment right posterior hippocampal activity was positively correlated with the path distance to the goal (Figure 3A, Table S4). This response was observed when including inverse efficiency scores (IES) in the model (see Supplemental Methods), to account for differences in reaction times and accuracy across events. This response was absent in the familiar environment and control routes, and there was also no parametric hippocampal response to IES on its own, underscoring the fact that it was not behavioural differences between environments that was driving the hippocampal response. Right posterior hippocampal activity was significantly more correlated with the distance to the goal in recent navigation routes than the other conditions combined, albeit at a threshold of p < 0.005 uncorrected (Figure 3B). Conducting the same analysis used for Travel Periods of excluding the events where the goal was a long distance away revealed that the significant correlation between activity and distance to the goal was dependent on these long distances (Figure S4B).

**Figure 3:**
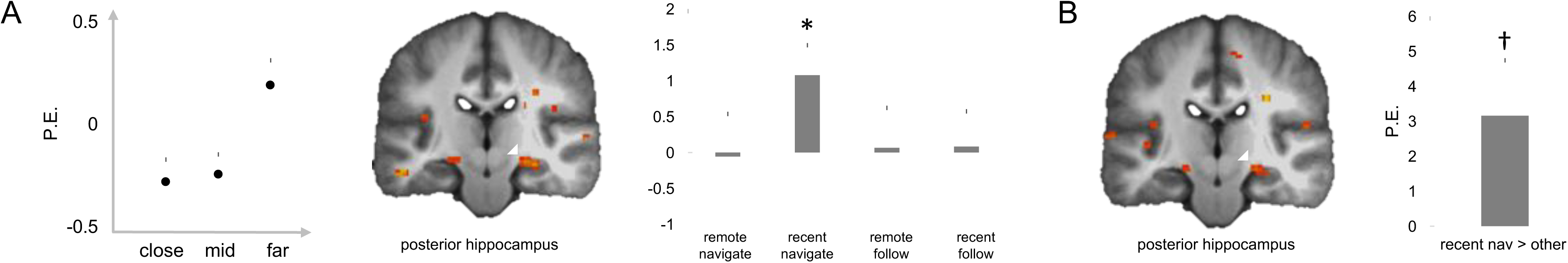
Path distance coding at Decision Points. A) At Decision Points, there was a significant positive correlation with path distance, such that there was higher BOLD activity in the right hippocampus when participants were further away from the goal location in recent environments. The effect plotted is corrected for IES (inverse efficiency), and is significant with and without this correction, thus underscoring that it is not a RT (or difficulty) effect. It also survives small-volume correction, including when ED (Euclidian distance) is added to the model. B) Brain activity when the GLM included weighted regressors for the effects seen in A. The contrast was −1 3 −1 −1, testing for an overall correlation with path distance during Decision Points, for the recent navigation condition. *=p<0.05 SVC, †p=0.005 u.c.

### Individuals who report more route planning in familiar environments have greater activity in their posterior hippocampus during the navigation of familiar environments

According to MTT/TTT and scene construction theory of hippocampal function (Moscovitch et al. 2005; Hassabis and Maguire, 2010), conscious mental simulation of future routes should engage the hippocampus no matter how familiar the environment may be. To test this prediction, we conducted a post-scan debriefing with a video replay of the navigation routes. At each Decision Point and New Goal Event the replay was paused and participants were asked whether they could recall planning their route to the goal. We found participants reported more route planning at events in recently learned environments than familiar environments (New Goal Events, Familiar: M=57%, SD=39; Recent: M=69%, SD=30; Decision Points, Familiar: M=8%, SD=10; Recent: M=22%, SD=20; paired t-test, both t>-3,p<0.007). Interestingly, in navigation routes in familiar environments, across participants, right posterior hippocampal activity was significantly correlated with the amount of reported planning at Decision Points (Figure 4A Table S4). This same measure of planning was also correlated with hippocampal activity during Travel Periods in familiar environments (Figure 4B, Table S4). We found performance accuracy at Decision Points did not correlate with hippocampal activity during navigation routes, either in the recently learned environment or the familiar environment (familiar and recent environments both: r<0.3, p>0.1), and that the amount of planning reported was not correlated with performance accuracy (familiar and recent environments both: r<-.06, p>0.1).

**Fig 4:**
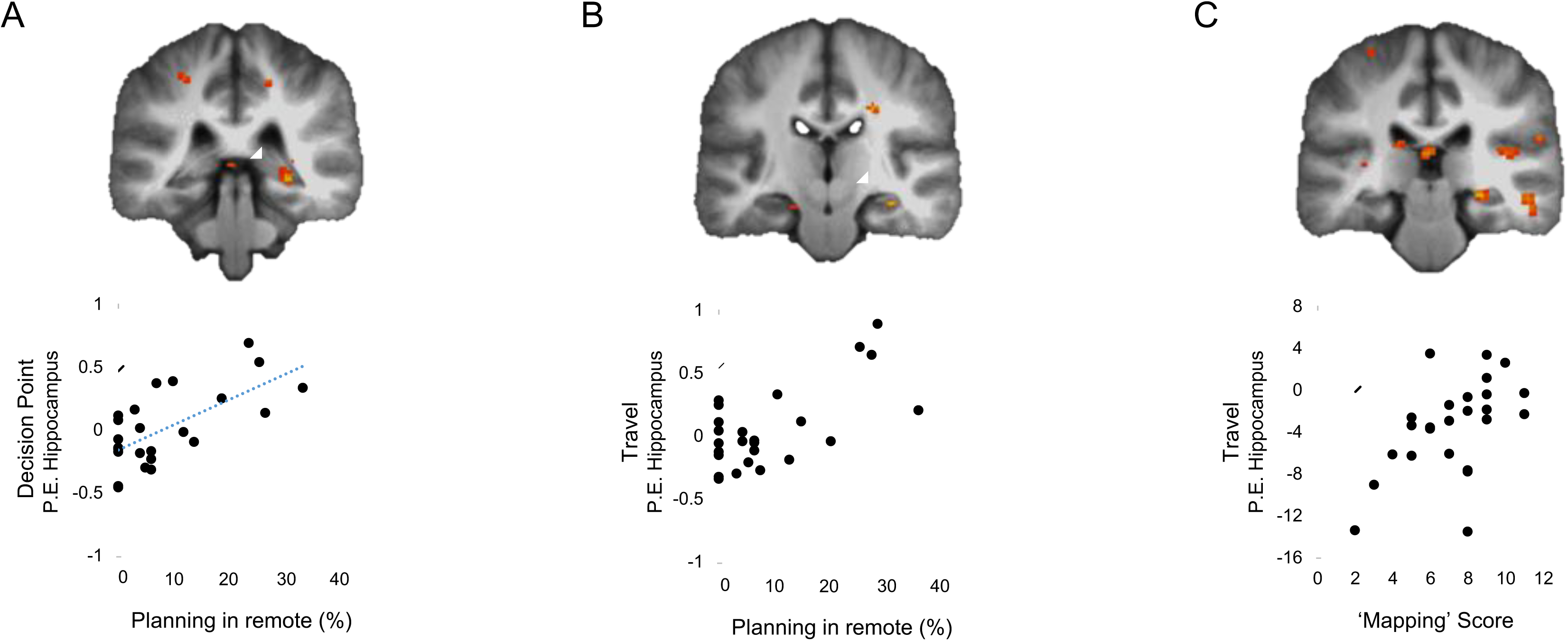
Right hippocampus active for people who plan and those who think in ‘map-space’. A) People who report planning more when at Decision points in remote environments show more right posterior hippocampal activity at Decision Points. B) People who report planning more when at Decision points in remote environments show more right posterior hippocampal activity during Travel. Both A&B (r>0.6,p<0.001). C) Top: People who report using more map-like strategies during navigation have more posterior hippocampal activity in relation to path distance during Travel in recent environments. Bottom: Plotted is the same effect, but extracting parameter estimates per person in the peak hippocampal voxel from Fig2A, and correlating it with mapping scores (r=0.53,p=0.007).

### Self-reported ‘map-based’ navigators have stronger correlations between hippocampal activity and the distance to the goal than ‘route-based’ navigators

Prior evidence indicates that strategy use for navigation impacts on the engagement of different brain regions for navigation of simulated environments (40, 41). To test whether this is true for navigation of real-world environments, participants completed a questionnaire probing navigational strategy use (see Supplemental Methods). The questionnaire determines the extent to which a person uses a map-based approach to navigation or a sequential landmark-based approach to navigation. Participants with higher map-based navigation scores had significantly stronger negative correlations between right posterior hippocampal activity and the distance to the goal during Travel Period in the recently learned environments (Figure 4C). We found no correlation between this self-reported strategy use and the amount of route planning (r<-0.18,p>0.1).

## Discussion

Using a virtual simulation of two London university campuses and fMRI we examined how, during navigation, the distance to the goal is represented by brain regions when the environment is highly familiar or recently learned. We found that in recently learned environments, the right posterior hippocampal activity is correlated with the distance to the goal, while in familiar environments distance to the goal was correlated with activity in the retrosplenial cortex. For recently learned environments, the more participants reported that they navigate via map-like strategies, the stronger the correlation between hippocampal activity and the distance to the goal. While overall we found that hippocampal activity was specific to a recently learned environment, participants who reported more route planning in a post-scan debrief showed increased hippocampal activity when navigating a familiar environment. These results help inform debates about memory consolidation, navigation systems of the brain, and the functional differentiation of the long-axis of the hippocampus.

### Systems consolidation of spatial representations

Our finding that hippocampal activity tracks the distance to the goal in recently learned, but not familiar environments, is consistent with standard consolidation models which argue that with the passage of time and repetition, information initially processed by the hippocampus becomes consolidated within neocortical structures (4, 42, 43). It is also consistent with evidence that lesions to the hippocampus lead to deficits in navigating new environments (44, 45), but not highly familiar environments (10, 17–21). As such, upon first inspection, our primary result would appear to conflict with the prediction from multiple trace theory (MTT) that detailed spatial processing, such as the distance to the goal, should always require the hippocampus no matter how familiar the environment (16). However, we speculated, in accord with TTT, that participants might alter how they navigate as the environment becomes more familiar and examined this with a post-scan debrief. Consistent with this prediction, we found that participants reported more route planning in the recently learned environment than the familiar environment. Moreover, we found that the more route planning a person reports in the familiar environment at decision points, the more hippocampal activity they express, which aligns well with the proposal in MTT/TTT that the hippocampus plays a continual role in supporting navigation when detailed processing is required (16). This finding is also broadly consistent with the argument that the hippocampal role in many tasks derives from a need to construct or re-construct scene representations (46). Our findings suggest that the amount of time spent in an environment, making it familiar, may not be the key determinant of the brain regions engaged, rather it is the shift in how an individual navigates the environment.

The brain regions responsible for processing consolidated spatial representations have remained somewhat mysterious (10). Here we found that the retrosplenial cortex codes spatial information in highly familiar environments rather than in recently learned environments. This finding is consistent with fMRI evidence that hippocampal activity declines with learning the layout of a virtual environment, but the activity in the retrosplenial cortex increases, tracking the new learning of stable spatial relationships in the environment (47–49). It is also consistent with reports of disorientation in highly familiar environments after retrosplenial lesions (10, 50, 51).

### Representation of the distance to the goal

Our finding that hippocampal activity was correlated with the distance to the goal in recently learned environment agrees with prior fMRI reports of similar coding (26, 27, 29, 30) and supports models which argue the hippocampus computes information about the future path to goal for navigational guidance (52–54). The observation that this was specific to navigation routes and not present in control routes is consistent with prior evidence (26) that such distance tracking is not automatic and requires goal-directed navigation. It is possible that the hippocampal activity correlated with distance to the goal relates to the pre-activation of place cells along a route to the goal, or ‘forward-sweeps’ of activity towards the goal (55). However, such responses may also relate to the recently discovered path distance coding neurons in the CA1 hippocampal region of flying bats, where each cell expresses activity at a certain distance from the goal (56). More cells are active at distances close to the goal, which is consistent with our observation of increased hippocampal activity with proximity to the goal during Travel Periods. At Decision Points we found the reverse function, activity increasing with distance to the goal, indicating there may be differences in the way goals and future paths are represented during the stages of navigation. Activity at Decision Points may reflect the retrieval demands of activating place cells representing the portions of space ahead along the route (Spiers and Barry, 2015), such as occurs during sharp-wave-ripple events (Pfieffer and Foster, 2013).

Consistent with our previous study exploring distance coding in real-world environments (26), we found the hippocampus coded distance to the goal with a positive and negative relationship at different stages of navigational journey (i.e., Decision Points and Travel Periods). However, we found the opposite pattern of responses to our previous study. While we find goal proximity coding during Travel Periods, our previous study reported goal proximity coding at Decision Points. Though it is difficult to determine what might give rise to this difference, three prominent factors may play a role. Compared to Howard et al (2014), the current study included much longer distances (maximum 1km vs 400m), fewer junctions on the routes, and more wide-open views on the streets. We found that down-sampling the data to exclude the longer distances substantially diminished the correlation between hippocampal activity and distance, whereas randomly down sampling had much less of an effect. This pattern is consistent with the system re-scaling the activity in relation to the dynamic range of the stimuli used, as has been found for value coding neurons in orbitofrontal cortex (57). Recent work indicates that turns within a route affect the representation of the distance within a space (58, 59), which may alter how the brain codes the distance to a goal within an environment. Notably, in a wide-open virtual environment, hippocampal activity was also found to be positively correlated with proximity to the goal during travel (Sherrill et al., 2013). Future research varying distance, turns, and visibility in the environment will be useful to further understand how neural systems encodes the spatial relationship to the future goal.

Computational models of navigation systems currently provide little specification of the implications of navigating a familiar environment compared with a recently learned environment. Our data indicate the retrosplenial cortex may play a key role in representing information about the distance to the goal in familiar environments. Because during Travel, hippocampus and retrosplenial cortex code distance in a different manner (negative and positive correlations with distance, respectively), it indicates that a model in which a simple switch from hippocampal cells coding the distance to retrosplenial cells coding the distance is not supported by our data. Rather MTT/TTT argue that once there is a change in neural representation, there also will be a change in functional or computational representation. It is possible that in a familiar environment, where there is less demand on route planning, the retrosplenial cortex plays a more prominent role in coding distance to the goal. Such a perspective is consistent with recent fMRI evidence of retrosplenial cortex positively correlated with the distance back to the start of a simple circular journey (25). It is also consistent with recent evidence of travel distance coding in rats during a route traversal task (60).

### Functional differentiation within the hippocampus

Similar to our previous study (26), we found a right posterior hippocampal focus for activity correlated with the distance to the goal. Prior neuropsychological research has consistently indicated the dominance of the right hemisphere in processing spatial information (45, 61, 62) and the posterior locus agrees with the view that posterior hippocampus processes fine detail, such as spatial metric information (39). Our data provide further support for this perspective in that the right posterior hippocampal activity is modulated by both the propensity for route planning in familiar environments and the extent to which a person reports using map-based strategies for navigation.

## Conclusion

In summary, our fMRI study provides the first comparison of active navigation of both a recently learned and a highly familiar environment. Our data supports models in which there is change in demand on brain regions with extended consolidation of the memories of an environment, with the hippocampus representing the distance to the goal in recently learned and the retrosplenial cortex in familiar environments. We also find support for the view that detailed spatial processing of an environment will involve the hippocampus even when recalling highly familiar environments (6, 16), and that this will entail the right posterior hippocampus (39, 62). Future research will be useful to determine how neuronal-level activity in the hippocampus and retrosplenial cortex may give rise to the fMRI signal dynamics reported here.

## Methods

### Participants

Students from the University College London (UCL) and Imperial College London campuses participated in this experiment. Recruitment involved selecting students who had been studying at either campus for a minimum of two years, and had little or no familiarity with the other university campus. This was assessed in a screening interview, in which participants had to label street names and landmarks on a blank map of the campuses. We collected twenty-six datasets, but one participant was excluded due to below chance performance during the fMRI session, resulting in the final sample of twenty-five subjects (12 UCL and 13 Imperial; mean age: 23 years, range: 20-26; 12 male [5 UCL,7 Imperial], 13 female [7 UCL, 6 Imperial]). Participants were administered two questionnaires regarding their navigation abilities and strategies (Santa Barbara Sense of Direction Scale [SBSDS] (63) and Navigational strategies questionnaire [NSQ], developed in Toronto by J.D.O. and J.R.).

All participants had normal or corrected to normal vision, reported no medical implant containing metal, no history of neurological or psychiatric condition, colour blindness, and did not suffer from claustrophobia. All participants gave written consent to participate to the study in accordance with UCL research ethics committee and the Birkbeck-UCL Centre for Neuroimaging (BUCNI) ethics committee. Participants were compensated with a minimum of £70 plus an additional £10 reward for their good performance during the scan.

### Training

The design of the experiment was based on the navigation task reported in Howard et al. (2014). There were, however, two campuses in which participants had to navigate: their native familiar campus (‘familiar’ environment) and the alternative new campus (‘recent’ environment). All participants needed to learn 10 goal locations, 18 streets and 8 start points in both environments. Participants were given training materials to practice for a week before the guided tour and the scanning session. Participants were trained on both the recent and familiar campuses in real life with a guided tour by an experimenter, with a strict set of rules, which were as follows: 1) Each road had to be walked past twice, in both directions, and each goal location had to be visited twice. 2) A probe for the name of each goal location was asked once before each visit (experimenter pointed in the direction of the nearest goal location before it became visible). 3) After arriving at each goal location, its name was read to the participant, and the direction of the start location was also given if the goal location was also a starting point. 4) On five occasions, participants were asked to point out the directions of two goal locations that they had visited twice. 5) The name of each street was asked twice, while the participant was not on it, before and after visiting it. 6) At the end of training, participants were asked about the directions of 10 goal locations and the names of the streets where they were located. The order in which participants were trained on the campuses was counterbalanced across participants and familiarity, and was done to ensure that the familiar campus was also recently visited in its entirety, thereby removing any confounding effects of just the recency of exposure (rather than the age of the memory itself). See Figure 1 for summary.

### Task Design

The task in the scanner was designed to simulate walking through the campuses, by using panorama images from Google Maps Street View (© Google 2014), to allow smooth and continuous navigation (developed by J.D.O). A large part of Imperial and a small part of UCL campuses were not mapped in Google Maps Street view, and these were substituted by panorama images taken by the experimenters (DMD Panorama, Dermandar S.A.L.). The photographs were taken every 6 meters. The latitude and longitude coordinates of each panorama image was extracted from Google Earth (© Google 2014) for precision. These images were then incorporated in the program filling the gaps left by Google Maps Street View.

Participants performed 16 routes in the scanner: 8 in the familiar and 8 in the recently learned campus. Half of these were ‘navigation’ blocks, i.e. participants had to actively navigate. In the other half, the ‘control’ condition, participants were led along the route and only had to make non-location based judgements. Each route began with a ‘New Goal Event’ to which they were required to navigate. In the navigation condition, participants were asked the general direction of this New Goal (‘Left or Right?’) and in the control condition, participants were asked ‘Can you buy a drink there?’. Following this screen, the program moved to the next panorama image, along the route. These will be referred to as ‘Travel Period’ events. When a junction was reached, participants were asked to choose which direction to go (‘Decision Point’: ‘Left’, ‘Right’, ‘Straight’ in navigation condition and this was given as an instruction with only one option in the control condition), after which the scene would pan into the next street (‘Turns’). As the routes were pre-determined, and the response the participant gave had no bearing on the actual trajectory, sometimes the ‘Turn’ events were in fact ‘Detours’, in which case a non-optimal path was taken. These events were infrequent, but were included to mimic real-life situations in which travel plans need to be updated. Please see Figure 1 for example task structure and timings. The total number (on average per subject) of each event type was as follows: Travel = 134, New Goal Events (NGE) = 46, Decision Points (DP) =107, Turns=76, Detour =33. Only about ¼ of these values were present per condition. Therefore, we will focus only on Travel periods and Decision Points, as they have a sufficient number of events (>20 per condition).

### Post-scan Debrief

Immediately after the scan there was a brief interview. All navigation routes that each participant was tested on were replayed in the same order as in the scanner. Participants were instructed to report what they remembered thinking during the navigation, not what they should have done, and to answer questions posed by the experimenter. At the start of each route they were asked “Were you oriented from the beginning?” – this was during the screen shown at the street entry. After that the experimenter pressed the play button. The navigation automatically paused whenever a New Goal Event appeared. Before and after each junction participants were told the responses made in the scanner and the experimenter would ask “Were you planning the route to the goal at this point during the scanning?”. At detours they were asked “Were you re-planning at this point?”. They were also asked if they were lost after detours. To this end, we acquired data at the following events (per familiar and recent campus): oriented, lost, and planning (at New Goal Events, Decision Points and Detours). Participants were also asked to report any salient memory at any point during navigation. All interviews were audio recorded.

### Spatial Parameters

Calculation of spatial parameters was done as in Howard et al., (2014) Javadi et al., (2017). In short, Path Distance (PD), Euclidian Distance (ED) and Egocentric Goal Direction (EGD) were extracted from the data. These parameters were then entered into fMRI analyses as regressors at all event types. Please see Table S1 for correlation between spatial parameters at each event type. The aim was to create routes where the spatial parameters were maximally decorrelated. However, due to the nature and layout of the campuses, there were limits on the flexibility of route design. Therefore, we entered path distance independently as a parametric regressor in our analyses, as this was the main focus of our task. However, we also checked our results when including ED along with PD, to establish the robustness of our findings, even with highly correlated regressors. Spatial parameter values were scaled between 0 and 1.

### fMRI Scanning & Preprocessing

Scanning was conducted at the Birkbeck-UCL Centre for Neuroimaging (BUCNI) using a 1.5 Tesla Siemens Avanto MRI scanner (Siemens Medical System, Erlangen, Germany) with a 32 channel head coil. Each experimental session lasted around 54 minutes and was separated in three parts (each of approximately 15-20 minutes). Approximately 980 functional scans were acquired per session (depending on routes taken), using a gradient-echo incremental EPI sequence (TR=3400ms, TE=85ms, TA=3.315s, flip angle=90°, 40 slices; slice thickness was 2mm with a gap of 1mm, TR=85ms, TE=50ms, slice tilt = 30°. The field of view was 192mm, and the matrix size was 64×64). The scan was a whole brain acquisition, with 40 slices. A T1-weighted high-resolution structural scan was acquired after the functional scans (TR=12ms, TE=5.6ms, 1×1×1mm^3^ resolution). Ear plugs were used for noise reduction, foam padding was used to secure the head in the scanner and minimize head movements. Stimuli were projected to the back screen, a mirror was attached to the head coil and adjusted for the subjects to see full screen.

All fMRI preprocessing and analysis was performed using SPM12 (Statistical Parametric Mapping, Wellcome Trust, London, UK). To achieve T1 equilibrium, the first six dummy volumes were discarded. During preprocessing, we used the SPM segment with 6 tissue classes to optimise normalisation. Otherwise, we used all default settings, and we performed slice timing correction. No participants had any abrupt motion changes over 4mm. Scanning was performed in 3 blocks, and as some events occurred rarely, we had to concatenate the fMRI data. We added a session regressor to indicate the change in scanning block.

### fMRI Analysis

For the fMRI analysis, we built multiple models based on a priori predictions from previous work (Howard et al, 2014). Please see Table S2 for a description of the models, events included and regression parameters (if applicable). Note for parametric modulation models, the event of interest was modelled with the corresponding spatial parameter regressors (i.e., Path Distance, Euclidian Distance, and Egocentric Goal Direction), but also included the other events in order to fully account for activity relating to stimulation. Additionally, we also included a task block regressor, which indicated whether the task was performed in a familiar or recent environment, and navigation or control. Only the implicit baseline (fixation period) of 17 seconds between routes was not included in the model. For example, when modelling Travel Periods, the model would include all Travel Period events in the 4 conditions (familiarity × navigation) + parametric modulators (pmods), in addition to the other events: DP, NGE, Detours, Turns, and Session, for each of the 4 conditions. Note that Travel Periods were defined as a single point in time while travelling down a route, and so was modelled as a punctate event using a stick function. Control models are described in detail in the Supplemental Results section. Small volume correction was done with defined anatomical masks (see Supplementary Methods for details).

## Acknowledgments

This work was supported by the Wellcome Trust (Grant 094850/Z/10/Z to HJS), James S. McDonnell Foundation (HJS), and Canadian Institute of Health Research (Grant # 49566 to MM and SR).

## Supplementary Methods

**Table S1:** Correlation between Event Types and raw Spatial Parameters.

| EVENT | PD-ED | PD-EGD | ED-EGD |
| --- | --- | --- | --- |
| **Decision Point** | 0.78** | 0.38** | 0.19* |
| **Travel** | 0.67** | 0.62** | 0.26** |

\*\*p<0.001; \*p<0.05, Bonferroni corrected; ~p<0.05

**Table S2:** Details of GLM parameters for the fMRI models

| MODEL | CONDITIONS | PMODS |
| --- | --- | --- |
| **CATEGORICAL- all events** | **Environment (2) x Nav (2)** | **IES\*** |
| **Travel** | **Environment (2) x Nav (2)** | **PD** |
| **Decision Point** | **Environment (2) x Nav (2)** | **PD, IES\*** |
| **CONTROL MODELS** |  |  |
| Travel | Environment (2) x Nav (2) | PD, ED |
| DP | Environment (2) x Nav (2) | PD, ED, IES* |
| Travel – long distances removed (25% of total) | Environment (2) x Nav (2) | PD |
| DP – long distances removed (25% of total ) | Environment (2) x Nav (2) | PD, IES* |
| Travel – random trials removed (25% of total ) | Environment (2) x Nav (2) | PD |
| DP – random trials removed (25% of total ) | Environment (2) x Nav (2) | PD, IES* |
| Travel – midpoints between events only | Environment (2) x Nav (2) | PD |
| Travel – New Street entry only | Environment (2) x Nav (2) | PD |
| Travel – Dead Ahead Goals only (each step modelled) | Environment (2) x Nav (2) | PD |
| Travel – few turns in segment (<3) | Environment (2) x Nav (2) | PD |
| Travel – many turns in segment (>2) | Environment (2) x Nav (2) | PD |
| Travel | Environment (2) x Nav (2) | Turns (# upcoming) |
| **ACROSS SUBJECTS MODELS** |  |  |
| Travel | Environment (2) x Nav only | DP planning scores |
| DP | Environment (2) x Nav only | DP planning scores |
| Travel PD | Recent Nav** | NSQ mapping scores |

\*Inverse efficiency: trial-by-trial RT/mean accuracy to control for differences in RT between familiar and recent condition during navigation (see behavioural results below)
\*\*Note that the GLM was run post-hoc after finding a significant correlation using extracted parameter estimates per person (in the peak hippocampal voxel for the Travel PD effect (recent navigate)) and mapping scores.

| EVENTS | DURATIONS |
| --- | --- |
| Decision Point | 5s |
| New Goal Event | 9s |
| Detour | 6s |
| Turns | 6s |
| Travel | 0s |
| Session | variable |

**Table S3:** Correlation between reaction times and spatial parameters across all subjects (r-values shown)

| OVERALL | PD | ED | EGD | PDxEGD<sup>†</sup> |
| --- | --- | --- | --- | --- |
| DP_RT | 0.31** | 0.23* | 0.05 | 0.27** |
| NGE_RT | 0.26~ | 0.02 | 0.19 | 0.27* |

| FAMILIAR | PD | ED | EGD | PDxEGD<sup>†</sup> |
| --- | --- | --- | --- | --- |
| DP_RT | 0.28** | 0.17~ | 0.02 | 0.26** |
| NGE_RT | 0.21 | 0.06 | 0.15 | 0.21 |

| RECENT | PD | ED | EGD | PDxEGD<sup>†</sup> |
| --- | --- | --- | --- | --- |
| DP_RT | 0.24** | 0.21* | 0.07 | 0.22* |
| NGE_RT | 0.29* | -0.02 | 0.23 | 0.33* |

\*\*sig. $p < 0.01$ , \*sig $p < 0.05$ , ~trend $p < 0.08$ , <sup>†</sup>to compare to Howard et al, 2014

## Supplementary Results

### Behaviour

In the pre-training assessment, we wanted to confirm that participants knew their own campus well, and were unfamiliar with the other campus. In the familiar (familiar) environment participants were on average 48% and 86% accurate on street names and landmarks, respectively. Conversely, for the recent campus, they were only 10% accurate on the street names, and 22% for landmarks. Thus, there was a significant effect of environment (F(1,24)=140.8,p<0.001), type of information probed (F(1,24)=48.9,p<0.001), and interaction (F(1,24)=14.8,p=0.001). Post-hoc paired t-test were also all significant (all t>3, p<0.004), underscoring the notion that participants did indeed have better knowledge about their own campus, and mainly its landmarks.

Our main interest was performance on the task performed during the scanning, which had a 2×2 factorial design (environment × navigation). We compared reaction times and accuracy at both New Goal Events (NGE) and Decision Points (DP), between familiar and recent environments, for both navigation and follow conditions using repeated-measures ANOVAs. At NGEs, there was a main effect of environment (F(1,24)=4.5,p=0.045), and a paired-t test revealed a trend for significantly lower RT in familiar navigate compared to recent navigate only (t=-1.9,p=0.064). For NGE accuracy, there was an overall effect of navigation (F(1,24)=41.8,p<0.001), with follow conditions having overall higher accuracy (as would be expected given participants were given the correct answer). For DP RTs, there was a main effect of navigation, environment and a significant interaction (all F>13.3, p=0.001). Paired-t tests confirmed that in the navigation condition, there was a significantly lower RT to DPs in familiar environments (t=-4.7, p=0.001). The same pattern was seen for DP accuracies. To control for potential differences in brain activation due to reaction time differences in the familiarity condition, we included a trial-by-trial inverse efficiency score (RT/mean accuracy) as a regressor when modelling the DPs.

For the debrief session, we compared responses to familiar and recent environments and found that although participants were equally oriented in both environments during NGEs (t=1.7, p>.1) and were only occasionally lost (less than 1%, but more so in the recent: t=-2.1, p=0.043), they did report more planning at NGEs, DPs, and Detours in the recent environment (all t>-2.9, p<0.006). We also found that participants who planned more at NGEs and DPs in familiar environments also planned more in recent ones (NGE: r>.88,p<0.001; DP: r=.6,p=0.001), and there was a correlation between how much one planned between NGEs and DPs in recent environments, (r=0.41,p=0.042), but not in familiar environments. In relation to the questionnaire measures, there was a trend towards a correlation between the NSQ & being lost in familiar environments (r=-.37, p=0.06), indicating that participants with higher scores on the questionnaire tended to be less lost. Relating performance in the scanner to debrief responses we found that NGE RT in familiar environments correlated with amount of planning at recent DPs (r=.46, p=0.02), such that when they were quicker to respond to goal, it meant less planning at decision points. Additionally, DP RT in recent environments correlated with amount of planning at DPs (both familiar and recent; >r=.42, p=0.03), such that less planning at DPs meant quicker responses. However, there was no correlation between planning and accuracy (all r<-.16, p>0.1).

We also looked at behaviour and its relation to spatial measures (for the navigation condition only), and found that both DP RTs in both environments, and NGE RTs in recent environments correlated significantly with PD (r>0.24, p<0.05), such that larger path distances incurred longer RTs. See Table S3 below for correlation coefficients.

Finally, we investigated the strategies questionnaires. Participants scored an average of 4.2 (range: 2.7-5.7) on the SBSDS. On the NSQ, they scored an average of 7.2 points (out of 14, where the maximum indicates only map-based navigation strategies). There was no correlation between the scores on these two questionnaires, between these scores and performance in the scanner, or to the amount of planning reported.

### Imaging

#### Categorical Effects

We explored global differences in brain activity, at each event of interest relating to the navigation vs follow condition, in both environments (see Figure S3). For familiar environments there was significantly greater activity in the precuneus for Travel, New Goal Events and Detours, and in the retrosplenial cortex for New Goal Events and Decision Points (Travel periods also showed this pattern but did not survive small volume correction [SVC]). New Goal Events also activated the visual cortex and intraparietal attention areas during navigation. The anterior cingulate region was also more active during navigation at Detours. This pattern of results is broadly consistent with the networks found during navigation in a similar paradigm reported by Howard et al (2014), however, our a-priori hypothesis was that this study would be more comparable to the ‘recent’ environment. Activity in recent environments in the current study were virtually absent when contrasting navigation vs follow conditions, perhaps reflecting engagement of similar networks, or continued encoding, when in a newly learned environment, regardless of the task demands.

**S1:**
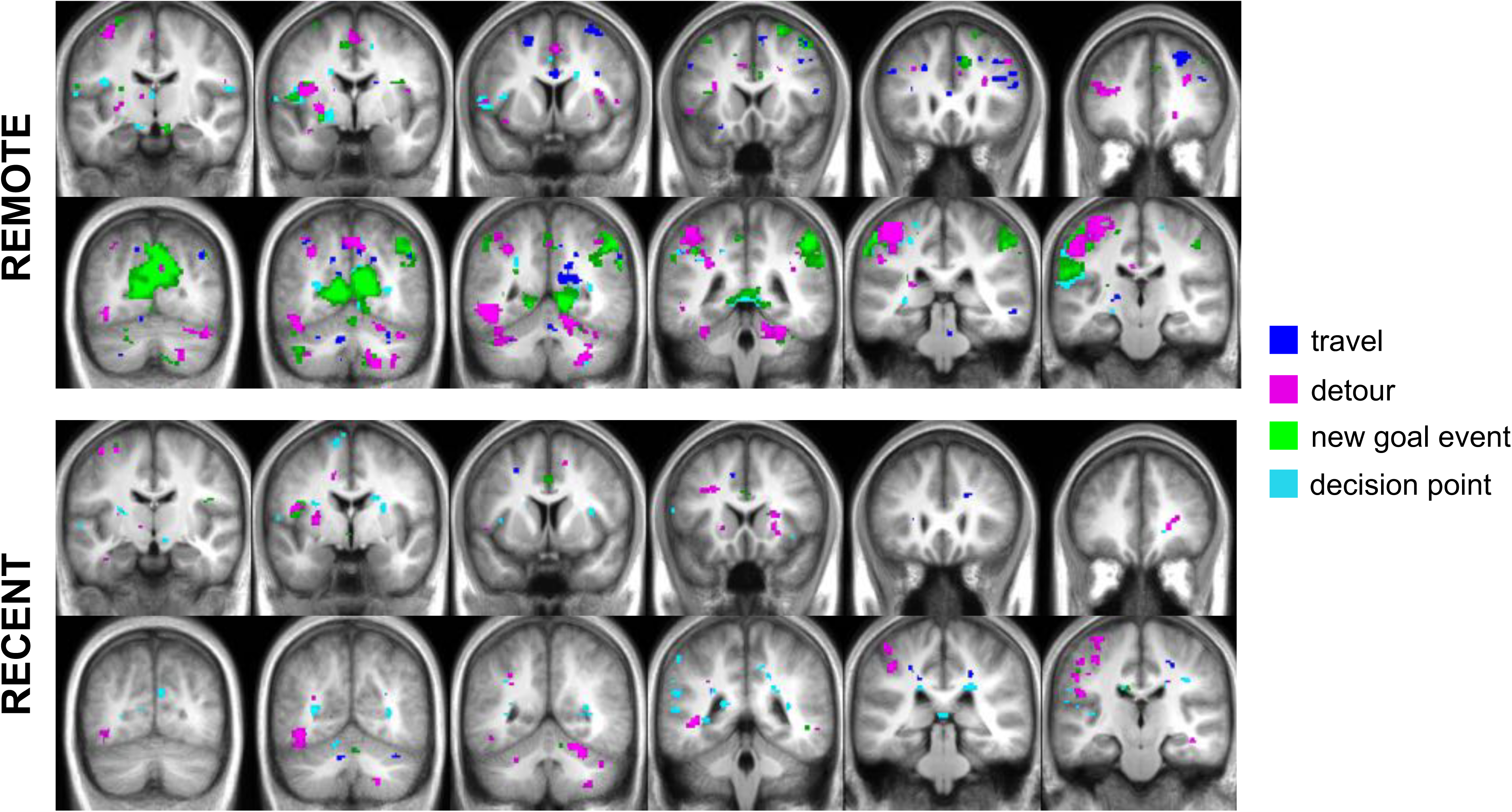
Categorical effects of all events for both familiar and recent environments (navigate > follow).

We were also interested how navigation in familiar vs recent environments may rely on different neural substrates. When looking at what areas were more active in the familiar environments (compared to recent, and only during navigation), we found again the precuneus during Detours and the retrosplenial cortex during Travel periods. Additionally the anterior cingulate cortex was more active at Decision Points. For the reverse contrast, the retrosplenial cortex showed increased activity during Detours. Activity at Decision Points was characterized by widespread temporal lobe and insular activation, as well as a more posterior to the ACC, dorso-medial PFC activation. More strikingly, when looking at overall session blocks of navigation in familiar vs recent, there was a clear and significant left hippocampal activation. All these contrasts are reported in Figures S2 and S3, as well as Table S4, and all effects reported are thresholded at p=0.001 uncorrected.

**S2:**
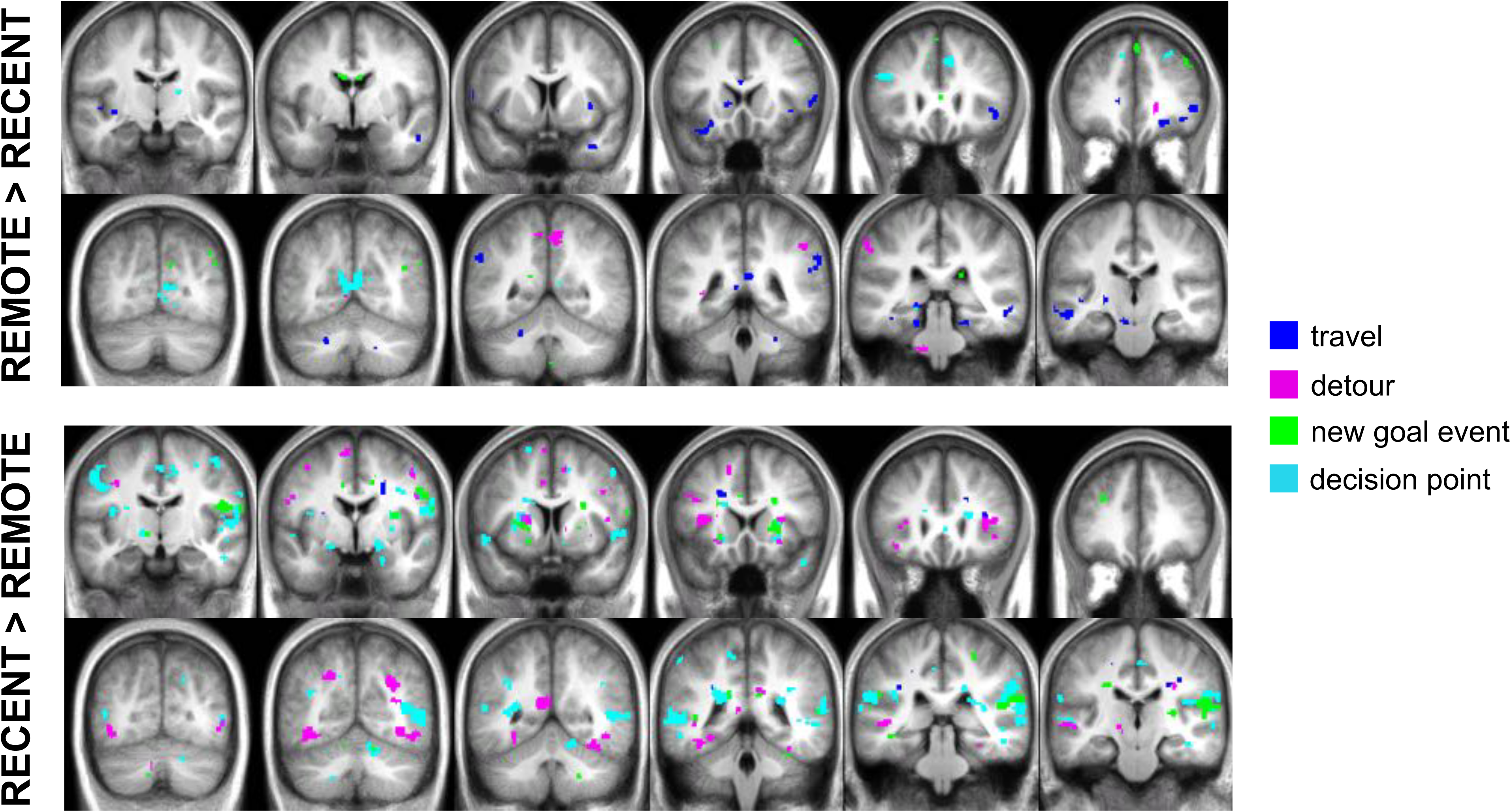
Categorical effects of all events for familiar > recent (navigation only)

**S3:**
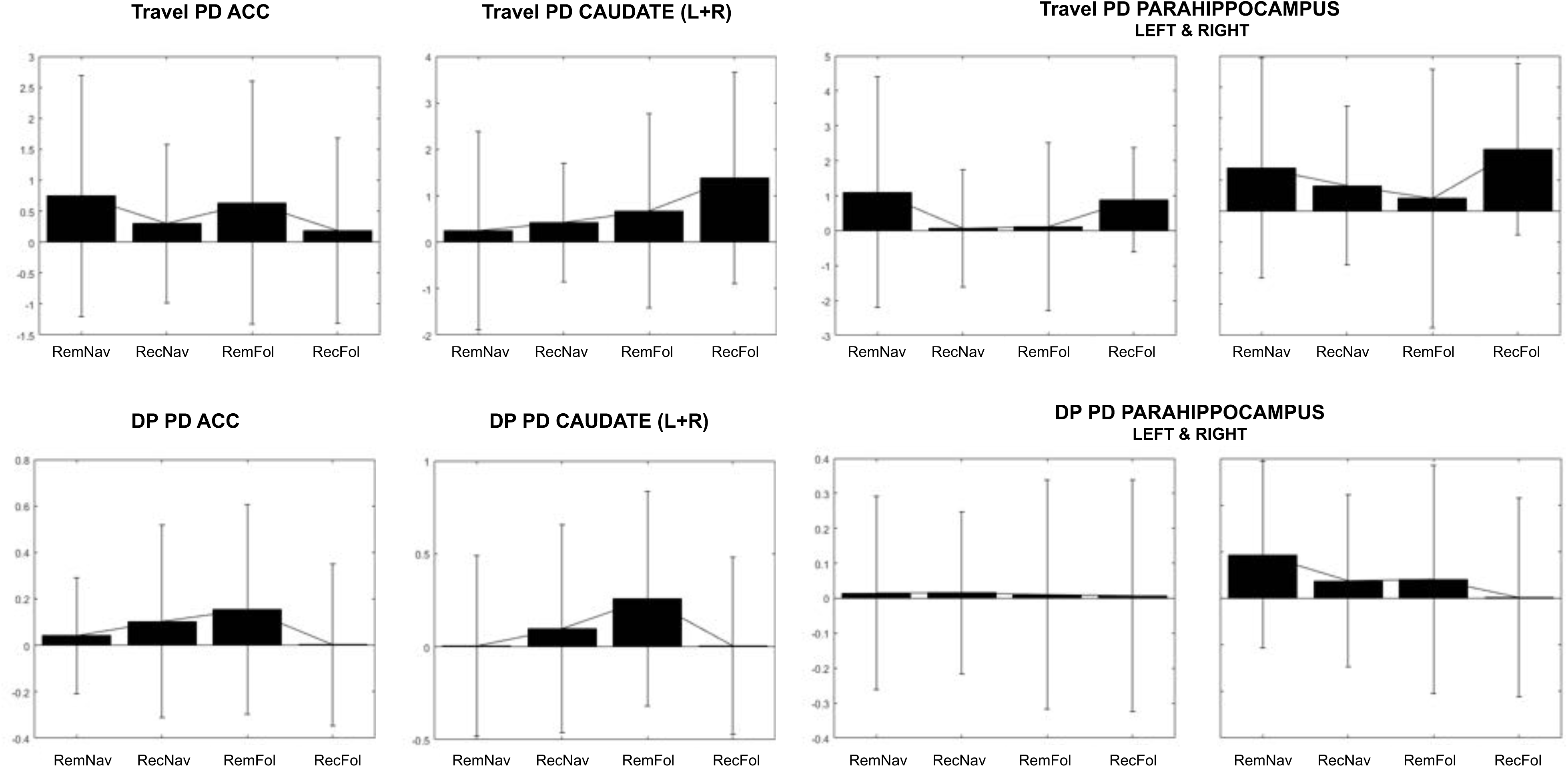
Parameter estimates from a-priori ROIs involved in navigation and memory. None of these areas seen at p=0.005 for any of the contrasts in Fig2/3, and no areas or conditions survive small volume correction.

**S4:**
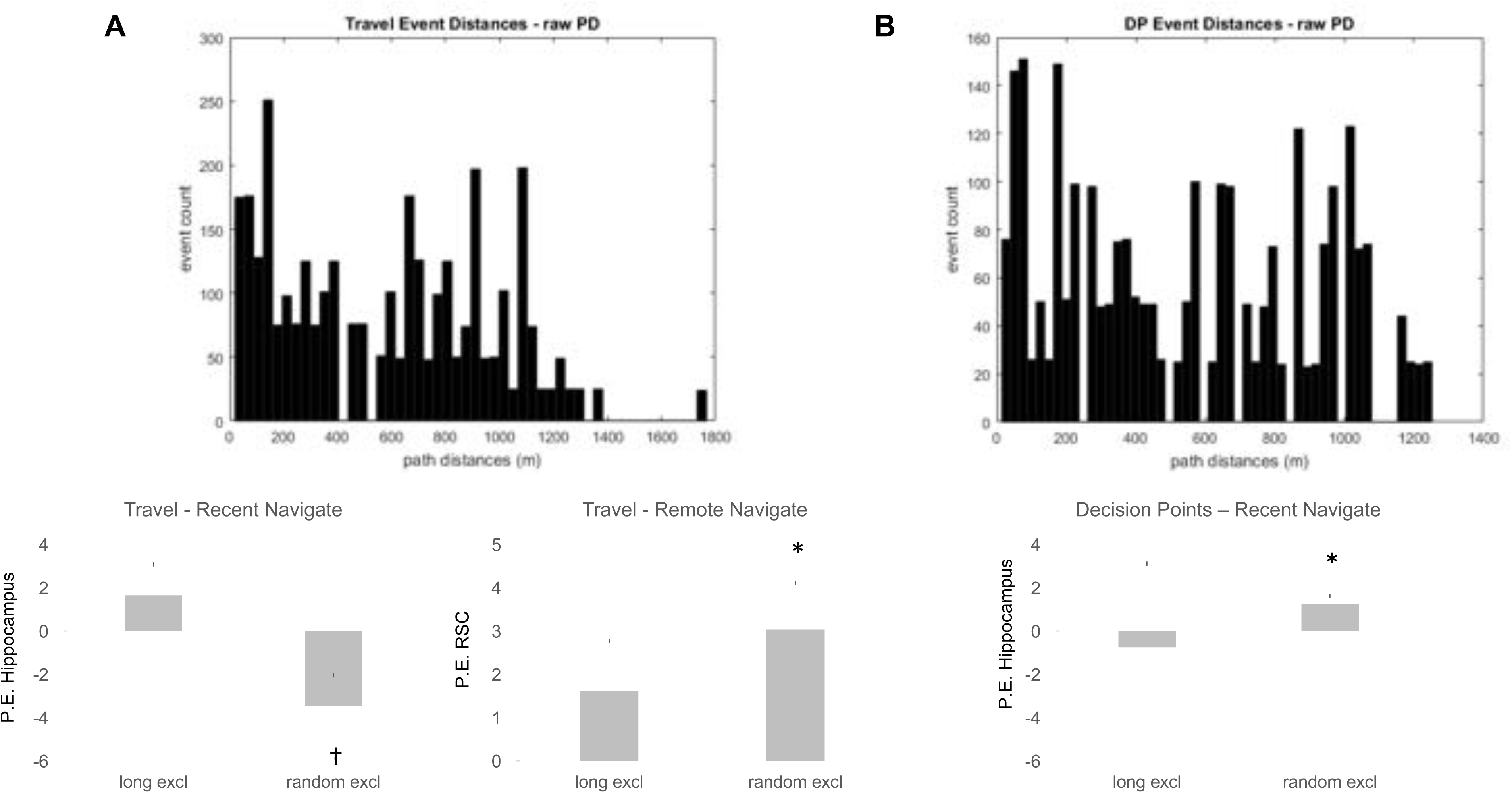
Correlations in Travel periods and Decision Points are driven primarily by long distances. A) Top: Distribution of path distances during Travel events. Bottom: Parameter estimates from the hippocampus and retrosplenial cortex for two models: 1) distances over 900m removed and 2) random trials removed subsampled to the same trial number as model 1. The effects with path distance are absent when large distances are removed. B) Top: Distribution of path distances during Decision Points events. Bottom: Parameter estimates from the hippocampus for two models as in Travel. *=p<0.05 SVC, †p<0.1 SVC

#### Control Analyses for effects of Path Distance

For all analyses regarding path distance we chose to focus on Travel Periods and Decision Points as they occurred most frequently, and contributed a minimum of 20 trials per condition.

We also ran control analyses, in which we included Euclidian distance (ED) in the models investigating parametric effects of path distance (PD). We replicated the PD results, emphasizing the robustness of these effects (see Table S4), despite highly correlated parametric regressors (see Table S1).

As our original Travel analysis included both travel midpoints as well as New Street Entry events, we also ran two additional models, which separated these events. When looking at Travel midpoints only, we replicated the right hippocampal effect in recent navigation, though this only survived a mid-hippocampal SVC (Table S4). We did not find the retrosplenial cortex in familiar navigation, but there was a positive correlation of PD in the precuneus and parietal-occipital sulcus. These effects were absent from the New Street Entry only models, emphasizing that the correlation with PD was specific to the Travel periods.

We also checked whether there was any evidence that the negative correlation with PD coding in the hippocampus was related to approaching the goal when it was straight ahead. We modeled Travel periods when this was the case, point-by-point, for segments that were at least 7 seconds long (minimum 2 TRs, which was 3.4 seconds). Two subjects did not have segments long enough (this depended on the routes they were given and some routes had shorter ‘goal approach’ segments) and were not included in the analysis. We did not find any evidence for coding in the hippocampus for recent navigation, which may be due to the reduced power (overall less samples), or perhaps that given the visualization of the task, the goal was not actually visible on each panorama image and as such it wasn’t as clear as it would have been if it were a continuous movie stream. Finally, we explored how the topology of the routes may have affected distance coding. We looked at Travel events that were in associated with either few or many upcoming turns. For Travel periods involving ‘few’ upcoming turns (<3), the hippocampal and retrosplenial PD effects, in recent and familiar environments, respectively, were replicated, but the ROIs did not survive statistical thresholding. Interestingly, there were no significant effects in recent environments for ‘many’ upcoming turns, however, in familiar environments there was a significant hippocampal cluster, positively correlated with PD. Retrosplenial cortex was also correlated with PD but this was not significant. When the environment is complex, encompassing many turns or fragments, the hippocampus may be involved even in familiar environments. However the lack of this effect in the recent environment precludes any conclusive suggestions as to how PD may be coded depending on environmental complexity when there is a strong need for planning ahead. In addition we also ran a model in which the number of upcoming turns was included as a parametric regressor for Travel (instead of PD), and found no effects. Please see Table S4 for z-scores and cluster sizes for significant effects. For small-volume corrections of the hippocampus and retrosplenial cortex, we used anatomical masks. The hippocampal ROI encompassed the mid and posterior right hippocampus (modified from Howard et al 2014).

#### Egocentric Goal Coding

Because of recent evidence of goal-related egocentric and proximity combined modulation of activity in the hippocampus (Howard et al., 2014; Sarel et al., 2017) we also explored whether there was any such modulation at Decision Points. We found the mid hippocampus was more active the further away, and less directly ahead the goal was, in recent environments. We also found evidence of medial superior parietal activity correlated with the egocentric direction to the goal during Travel Period in the familiar environment, broadly consistent with prior findings (26, 28). Note egocentric goal direction was also highly correlated with path distance (Table S1; see TS4 for details of activation).

## References

1. Mcgaugh JL (2000) Memory — a Century of Consolidation. 287(January):248–252.

2. Wang S-H, Morris RGM (2010) Hippocampal-neocortical interactions in memory formation, consolidation, and reconsolidation. Annu Rev Psychol 61(November):49–79, C1-4.

3. Eichenbaum H, Yonelinas AP, Ranganath C (2007) The medial temporal lobe and recognition memory. Annu Rev Neurosci 30:123–52.

4. Frankland PW, Bontempi B (2005) The organization of recent and remote memories. Nat Rev Neurosci 6(2):119–30.

5. Squire LR, Wixted JT (2010) The Cognitive Neuroscience of Human Memory Since H.M. Annu Rev Neurosci (April):259–288.

6. Moscovitch M, Cabeza R, Winocur G, Nadel L (2016) Episodic memory and beyond: The hippocampus and neocortex in transformation. Annu Rev Psychol 67(1):105–134.

7. Tulving E, Markowitsch HJ (1998) Episodic and Declarative Memory: Role of the Hippocampus. Hippocampus 8:198–204.

8. Rosenbaum RS, Winocur G, Moscovitch M (2001) New views on old memories: re-evaluating the role of the hippocampal complex. 127:183–197.

9. Winocur G, Moscovitch M, Bontempi B (2010) Memory formation and long-term retention in humans and animals: convergence towards a transformation account of hippocampal-neocortical interactions. Neuropsychologia 48(8):2339–2356.

10. Spiers HJ, Maguire EA (2007) The neuroscience of remote spatial memory: A tale of two cities. Neuroscience 149(1):7–27.

11. Squire LR (1992) Memory and the Hippocampus: A Synthesis From Findings With Rats, Monkeys, and Humans. Psychol Rev 99(2):195–231.

12. Squire LR, Zola-Morgan S (1998) Episodic Memory, Semantic Memory, and Amnesia. Hippocampus 8:205–211.

13. Winocur G, Moscovitch M (2011) Memory Transformation and Systems Consolidation. J Int Neuropsychol Soc 17(5):766–780.

14. Nadel L, Moscovitch M (1997) Memory consolidation and the hippocampal complex. Curr Opin Neurobiol 7:217–227.

15. Vargha-khadem F, Gadian DG, Mishkin M (2001) Dissociations in cognitive memory: the syndrome of developmental amnesia. Philos Trans R Soc Lond B Biol Sci 356(1413):1435–40.

16. Moscovitch M, et al. (2005) Functional neuroanatomy of remote episodic, semantic and spatial memory: a unified account based on multiple trace theory. J Anat 207(1):35–66.

17. Teng E, Squire LR (1999) Memory for places learned long ago is intact after hippocampal damage. Nature 400(6745):675–677.

18. Maguire EA, Nannery R, Spiers HJ (2006) Navigation around London by a taxi driver with bilateral hippocampal lesions. Brain 129(11):2894–2907.

19. Rosenbaum RS, et al. (2000) Remote spatial memory in an amnesic person with extensive bilateral hippocampal lesions. Nat Neurosci 3(10):1044–1048.

20. Herdman KA, Calarco N, Moscovitch M, Hirshhorn M, Rosenbaum RS (2015) Impoverished descriptions of familiar routes in three cases of hippocampal/medial temporal lobe amnesia. Cortex 71:248–263.

21. Rosenbaum RS, Gao F, Richards B, Black SE, Moscovitch M (2005) “‘Where to?’” Remote Memory for Spatial Relations and Landmark Identity in Former Taxi Drivers with Alzheimer’s Disease and Encephalitis. J Cogn Neurosci 17(3):446–462.

22. Spiers HJ, Maguire EA (2006) Thoughts, behaviour, and brain dynamics during navigation in the real world. Neuroimage 31(4):1826–1840.

23. Rosenbaum RS, Ziegler M, Winocur G, Grady CL, Moscovitch M (2004) “I have often walked down this street before”: fMRI studies on the hippocampus and other structures during mental navigation of an old environment. Hippocampus 14(7):826–835.

24. Hirshhorn M, Grady C, Rosenbaum RS, Winocur G, Moscovitch M (2012) The hippocampus is involved in mental navigation for a recently learned, but not a highly familiar environment: A longitudinal fMRI study. Hippocampus 22(4):842–852.

25. Chrastil ER, Sherrill KR, Hasselmo ME, Stern CE (2015) There and Back Again: Hippocampus and Retrosplenial Cortex Track Homing Distance during Human Path Integration. J Neurosci 35(46):15442–15452.

26. Howard LR, et al. (2014) The hippocampus and entorhinal cortex encode the path and Euclidean distances to goals during navigation. Curr Biol 24(12):1331–40.

27. Sherrill KR, et al. (2013) Hippocampus and retrosplenial cortex combine path integration signals for successful navigation. J Neurosci 33(49):19304–13.

28. Spiers HJ, Maguire EA (2007) A Navigational Guidance System in the Human Brain. Hippocampus 0(17):618–626.

29. Viard A, Doeller CF, Hartley T, Bird CM, Burgess N (2011) Anterior Hippocampus and Goal-Directed Spatial Decision Making. J Neurosci 31(12):4613–4621.

30. Balaguer J, Spiers HJ, Hassabis D, Summerfield C (2016) Neural Mechanisms of Hierarchical Planning in a Virtual Subway Network. Neuron 90(4):893–903.

31. Morgan LK, MacEvoy SP, Aguirre GK, Epstein RA (2011) Distances between Real-World Locations Are Represented in the Human Hippocampus. J Neurosci 31(4):1238–1245.

32. Takahashi N, Kawamura M, Shiota J, Kasahata N, Hirayama K (1997) Pure topographic disorientation due to right retrosplenial lesion. Neurology 49(2):464–469.

33. Epstein RA, Parker WE, Feiler AM (2007) Where am I now? Distinct roles for parahippocampal and retrosplenial cortices in place recognition. J Neurosci 27(23):6141–6149.

34. Bohbot VD, et al. (2015) Role of the parahippocampal cortex in memory for the configuration but not the identity of objects: converging evidence from patients with selective thermal lesions and fMRI. Front Hum Neurosci 9(August):431.

35. Epstein RA (2008) Parahippocampal and retrosplenial contributions to human spatial navigation. Trends Cogn Sci 12(10):388–96.

36. Ranganath C, Ritchey M (2012) Two cortical systems for memory-guided behaviour. Nat Rev Neurosci 13(10):713–726.

37. Miller a. MP, Vedder LC, Law LM, Smith DM (2014) Cues, context, and long-term memory: the role of the retrosplenial cortex in spatial cognition. Front Hum Neurosci 8(August):1–15.

38. Teixeira CM, Pomedli SR, Maei HR, Kee N, Frankland PW (2006) Involvement of the anterior cingulate cortex in the expression of remote spatial memory. J Neurosci 26(29):7555–7564.

39. Poppenk J, Evensmoen HR, Moscovitch M, Nadel L (2013) Long-axis specialization of the human hippocampus. Trends Cogn Sci:1–11.

40. Iaria G, Petrides M, Dagher A, Pike B, Bohbot VD (2003) Cognitive strategies dependent on the hippocampus and caudate nucleus in human navigation: variability and change with practice. J Neurosci 23(13):5945–5952.

41. Iglói K, Doeller CF, Berthoz A, Rondi-Reig L, Burgess N (2010) Lateralized human hippocampal activity predicts navigation based on sequence or place memory. Proc Natl Acad Sci 107(32):14466–14471.

42. Marr (1971) Simple memory: a theory for achicortex. Philos Trans R Soc London 262(841):23–81.

43. Alvarez P, Squire LR (1994) Memory consolidation and the medial temporal lobe: A simple network model. 91(July):7041–7045.

44. Spiers HJ, et al. (2001) Unilateral temporal lobectomy patients show lateralized topographical and episodic memory deficits in a virtual town. Brain 124(Pt 12):2476–89.

45. Spiers HJ, Maguire EA, Burgess N (2001) Hippocampal amnesia. Neurocase 7(5):357–82.

46. Hassabis D, Maguire EA (2009) The construction system of the brain. Philos Trans R Soc Lond B Biol Sci 364(1521):1263–71.

47. Wolbers T, Büchel C (2005) Dissociable retrosplenial and hippocampal contributions to successful formation of survey representations. J Neurosci 25(13):3333–40.

48. Auger SD, Mullally SL, Maguire EA (2012) Retrosplenial cortex codes for permanent landmarks. PLoS One 7(8):e43620.

49. Auger SD, Zeidman P, Maguire EA (2015) A central role for the retrosplenial cortex in de novo environmental learning. Elife 4(AUGUST2015):1–26.

50. Maguire EA, Vargha-khadem F, Mishkin M (2001) The effects of bilateral hippocampal damage on fMRI regional activations and interactions during memory retrieval. Brain 124:1156–1170.

51. Aguirre GK, D’Esposito M (1999) Topographical disorientation: A synthesis and taxonomy. Brain 122(9):1613–1628.

52. Erdem UM, Hasselmo M (2012) A goal-directed spatial navigation model using forward trajectory planning based on grid cells. Eur J Neurosci 35(6):916–931.

53. Penny WD, Zeidman P, Burgess N (2013) Forward and Backward Inference in Spatial Cognition. PLoS Comput Biol 9(12). doi:10.1371/journal.pcbi.1003383.

54. Bush D, Barry C, Manson D, Burgess N (2015) Using Grid Cells for Navigation. Neuron 87(3):507–520.

55. Spiers HJ, Barry C (2015) Neural systems supporting navigation. Curr Opin Behav Sci 1:47–55.

56. Sarel A, Finkelstein A, Las L, Ulanovsky N (2017) Vectorial representation of spatial goals in the hippocampus of bats. Science (80-) 180(January):176–180.

57. Padoa-Schioppa C (2009) Range-Adapting Representation of Economic Value in the Orbitofrontal Cortex. J Neurosci 29(44):14004–14014.

58. Bonasia K, Blommesteyn J, Moscovitch M (2016) Memory and navigation: Compression of space varies with route length and turns. Hippocampus 26(1):9–12.

59. Brunec IK, Javadi A, Zisch FEL, Spiers HJ (2017) Contracted time and expanded space: The impact of circumnavigation on judgements of space and time. Cognition 166:425–432.

60. Alexander AS, Nitz DA (2017) Spatially Periodic Activation Patterns of Retrosplenial Cortex Encode Route Sub-spaces and Distance Traveled. Curr Biol 27(11):1551–1560.e4.

61. Burgess N, Maguire EA, Spiers HJ, O’Keefe J (2001) A temporoparietal and prefrontal network for retrieving the spatial context of lifelike events. Neuroimage 14(2):439–53.

62. Burgess N, Maguire EA, O’Keefe J (2002) The human hippocampus and spatial and episodic memory. Neuron 35(4):625–41.

63. Hegarty, M., Richardson, A. E., Montello, D. R., Lovelace, K., & Subbiah I (2002) Development of a self-report measure of environmental spatial ability. Intelligence.

64. Javadi A-H, et al. (2017) Hippocampal and prefrontal processing of network topology to simulate the future. Nat Commun in press:1–11.

